# Synergy between c-di-GMP and quorum-sensing signaling in *Vibrio cholerae* biofilm morphogenesis

**DOI:** 10.1101/2022.06.27.497871

**Authors:** Jojo A. Prentice, Andrew A. Bridges, Bonnie L. Bassler

## Abstract

Transitions between individual and communal lifestyles allow bacteria to adapt to changing environments. Bacteria must integrate information encoded in multiple sensory cues to appropriately undertake these transitions. Here, we investigate how two prevalent sensory inputs converge on biofilm morphogenesis: quorum sensing, which endows bacteria with the ability to communicate and coordinate group behaviors, and second messenger c-di-GMP signaling, which allows bacteria to detect and respond to environmental stimuli. We use *Vibrio cholerae* as our model system, the autoinducer AI-2 to modulate quorum sensing, and the polyamine norspermidine to modulate NspS-MbaA-mediated c-di-GMP production. Individually, AI-2 and norspermidine drive opposing biofilm phenotypes, with AI-2 repressing and norspermidine inducing biofilm formation. Surprisingly, however, when AI-2 and norspermidine are simultaneously detected, they act synergistically to elevate biofilm formation. We show that this effect is caused by quorum-sensing-mediated activation of *nspS*-*mbaA* expression, which increases the levels of NspS and MbaA, and in turn, c-di-GMP biosynthesis, in response to norspermidine. Increased MbaA-synthesized c-di-GMP activates the VpsR transcription factor, driving elevated expression of genes encoding key biofilm matrix components. Thus, in the context of biofilm formation in *V. cholerae*, quorum-sensing regulation of c-di-GMP-metabolizing receptor levels connects changes in cell population density to detection of environmental stimuli.

**Importance:** The development of multicellular communities, known as biofilms, facilitates beneficial functions of gut microbiome bacteria and makes bacterial pathogens recalcitrant to treatment. Understanding how bacteria regulate the biofilm lifecycle is fundamental to biofilm control in industrial processes and in medicine. Here, we demonstrate how two major sensory inputs – quorum-sensing communication and second messenger c-di-GMP signaling – jointly regulate biofilm morphogenesis in the global pathogen *Vibrio cholerae*. We characterize the mechanism underlying a surprising synergy between quorum-sensing and c-di-GMP signaling in controlling biofilm development. Thus, the work connects changes in cell population density to detection of environmental stimuli in a pathogen of clinical significance.

## Introduction

Bacteria often integrate multiple sensory cues into the control of behaviors including the formation of biofilms – surface-associated bacterial communities encapsulated in self-produced extracellular matrices (1). The biofilm lifestyle confers advantages to constituent members, including protection against antibiotics, predation, and shear stress (2–4). Indeed, biofilms are a predominant form of bacterial life in the environment, in industrial processes, and in disease (5).

In the global pathogen and model biofilm-forming bacterium *Vibrio cholerae*, two well-studied sensory inputs control the biofilm lifecycle. The first is quorum sensing: a cell-cell communication process that orchestrates collective behaviors (6). Quorum sensing relies on the production, release, and group-wide detection of extracellular signaling molecules called autoinducers (7). *V. cholerae* possesses five quorum-sensing autoinducer-receptor pairs, two of which are key to the present work, diagrammed in Fig. 1 (8). At low cell density, the autoinducer receptors CqsS and LuxPQ are unliganded and function as kinases, channeling phosphate to the response regulator LuxO (9,10). LuxO∼P indirectly represses the gene encoding the high cell density master regulator HapR (11,12). HapR represses expression of the vibrio polysaccharide biosynthetic genes (*vpsI* and *vpsII* operons), *vpsT*, encoding a transcriptional activator of the *vpsI* and *vpsII* operons, and *rbmA* and *rbmC-E*, encoding biofilm matrix proteins. Thus, in the low cell density quorum-sensing regime, when *hapR* is repressed, VPS and biofilm matrix protein levels are high, and *V. cholerae* forms biofilms (13). At high cell density, cholerae autoinducer-1 (CAI-1) and autoinducer-2 (AI-2) accumulate and bind CqsS and LuxPQ respectively, converting them from kinases to phosphatases. Phosphate is stripped from LuxO, which inactivates it (9,10). As a result, HapR is produced, it suppresses biofilm formation, and biofilm dispersal occurs (Fig. 1) (12).

**Fig. 1.**
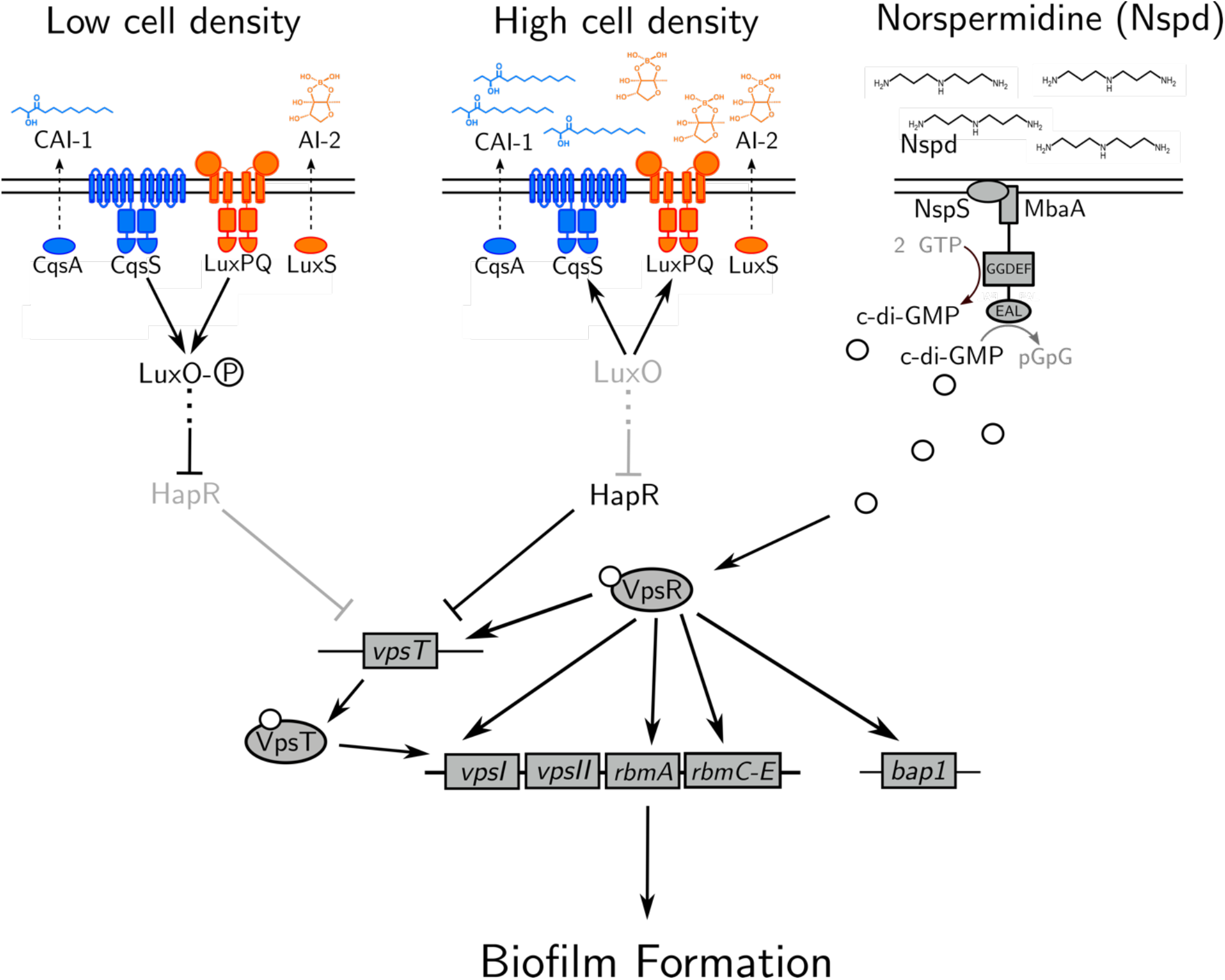
Model showing the contributions of quorum-sensing and norspermidine signaling to biofilm gene expression. See text for details. The P in the circle represents phosphate. White circles represent c-di-GMP. Norspermidine, Nspd.

The second major regulator of the *V. cholerae* biofilm lifecycle is the second messenger molecule cyclic diguanylate (c-di-GMP). c-di-GMP is produced and degraded by enzymes containing diguanylate cyclase and/or phosphodiesterase activities, respectively. These activities are commonly modulated by environmental stimuli including light, temperature, amino acids, oxygen, and polyamines (14–18). High intracellular c-di-GMP levels drive biofilm formation via binding to and activation of the VpsT and VpsR transcription factors. VpsT-c-di-GMP and VpsR-c-di-GMP both activate expression of the *vpsI* and *vpsII* operons, and additionally, VpsR-c-di-GMP activates expression of *rbmA, rbmC-E*, and *bap1*. By contrast, when cytoplasmic c-di-GMP levels are low, biofilm formation is repressed, favoring the motile state (Fig. 1) (18,19). Thus, in *V. cholerae*, the low cell density quorum-sensing regime and high levels of cytoplasmic c-di-GMP each promote biofilm formation, whereas the high cell density quorum-sensing regime and low levels of cytoplasmic c-di-GMP each repress biofilm formation. Attempts to knit together the *V. cholerae* quorum-sensing and c-di-GMP pathways have revealed two key findings: first, high cytoplasmic c-di-GMP concentrations can override negative quorum-sensing regulation of biofilm genes (21,22). Second, the high cell density quorum-sensing regime activates the expression of genes encoding over a dozen diguanylate cyclases and phosphodiesterases, while repressing only a few genes encoding such enzymes (8,21).

Here, we investigate the integration of quorum-sensing and c-di-GMP information in *V. cholerae* biofilm morphogenesis, from ligand detection to population-scale biofilm changes. We use exogenous administration of the AI-2 autoinducer to modulate quorum-sensing activity and we use administration of the polyamine norspermidine to control the activity of the NspS-MbaA c-di-GMP-metabolizing circuit (Fig. 1) (17). We find that as expected, quorum sensing represses biofilm formation in the absence of NspS-MbaA detection of norspermidine. However, surprisingly, quorum sensing promotes biofilm formation when MbaA-mediated c-di-GMP synthesis is stimulated by norspermidine supplementation. We show that this positive quorum-sensing effect occurs because at high cell density, HapR activates *nspS*-*mbaA* expression, which drives increased NspS and MbaA production and consequently, increased c-di-GMP production when the norspermidine ligand is present. The increased c-di-GMP activates VpsR, which in turn, activates *rbmA* matrix gene expression, resulting in the formation of larger and denser biofilms. We propose a model in which quorum sensing represses biofilms, but also primes the bacterial population to optimally respond to environmental stimuli that foster c-di-GMP production. Our findings reveal a new mechanism by which *V. cholerae* modulates its biofilm lifecycle and, moreover, they show that quorum sensing does not strictly repress *V. cholerae* biofilm formation.

## Results

### Quorum sensing promotes norspermidine-mediated biofilm formation

To investigate how *V. cholerae* integrates information from c-di-GMP and quorum-sensing signaling into the control of the biofilm lifecycle, we measured biofilm phenotypes across quorum-sensing and c-di-GMP signaling regimes. To simplify the regulation of quorum sensing, we used a *V. cholerae* strain harboring only a single quorum-sensing receptor that controls LuxO phosphorylation – the AI-2 receptor LuxPQ. Moreover, we deleted the AI-2 synthase *luxS* so that quorum sensing is exclusively controlled through exogenous administration of AI-2. We refer to this strain as the “AI-2-responsive strain.” First, we measured biofilm biomass accumulation over time in the AI-2-responsive strain in the absence of AI-2 (i.e., in the low cell density quorum-sensing regime, Fig. 2A,B). Consistent with previous findings, in this signaling regime, biofilms formed (Fig. 2A) (13). Addition of saturating AI-2 (i.e., to achieve the high cell density quorum-sensing regime, Fig. 2A,B) prevented the AI-2-responsive strain from forming biofilms, again, consistent with previous findings (13,21,23). To investigate how changes in c-di-GMP affect biofilm formation in the low and high cell density quorum-sensing regimes, we provided exogenous norspermidine to drive c-di-GMP production. Norspermidine had only a modest effect on peak biofilm biomass when the AI-2-responsive *V. cholerae* strain was in the low cell density quorum-sensing regime, whereas, surprisingly, norspermidine drove dramatically increased biofilm formation when the strain was in the high cell density quorum-sensing regime (Fig. 2A,B). These results were independent of the specific autoinducer-receptor pair used to stimulate quorum sensing in *V. cholerae*, as we likewise modulated quorum sensing in a CAI-1-responsive strain and obtained analogous results (Fig. S1A). Moreover, the observed increase in biofilm biomass in the high cell density quorum-sensing and high norspermidine regime did not occur when *mbaA* was deleted (Fig. S1B). This result shows that changes in biofilm formation are mediated by the known polyamine-sensing NspS-MbaA pathway. Thus, although quorum sensing and norspermidine independently drive opposing biofilm phenotypes, with quorum-sensing repressing and c-di-GMP promoting biofilm formation, together, they function synergistically to foster biofilm formation.

**Fig. 2.**
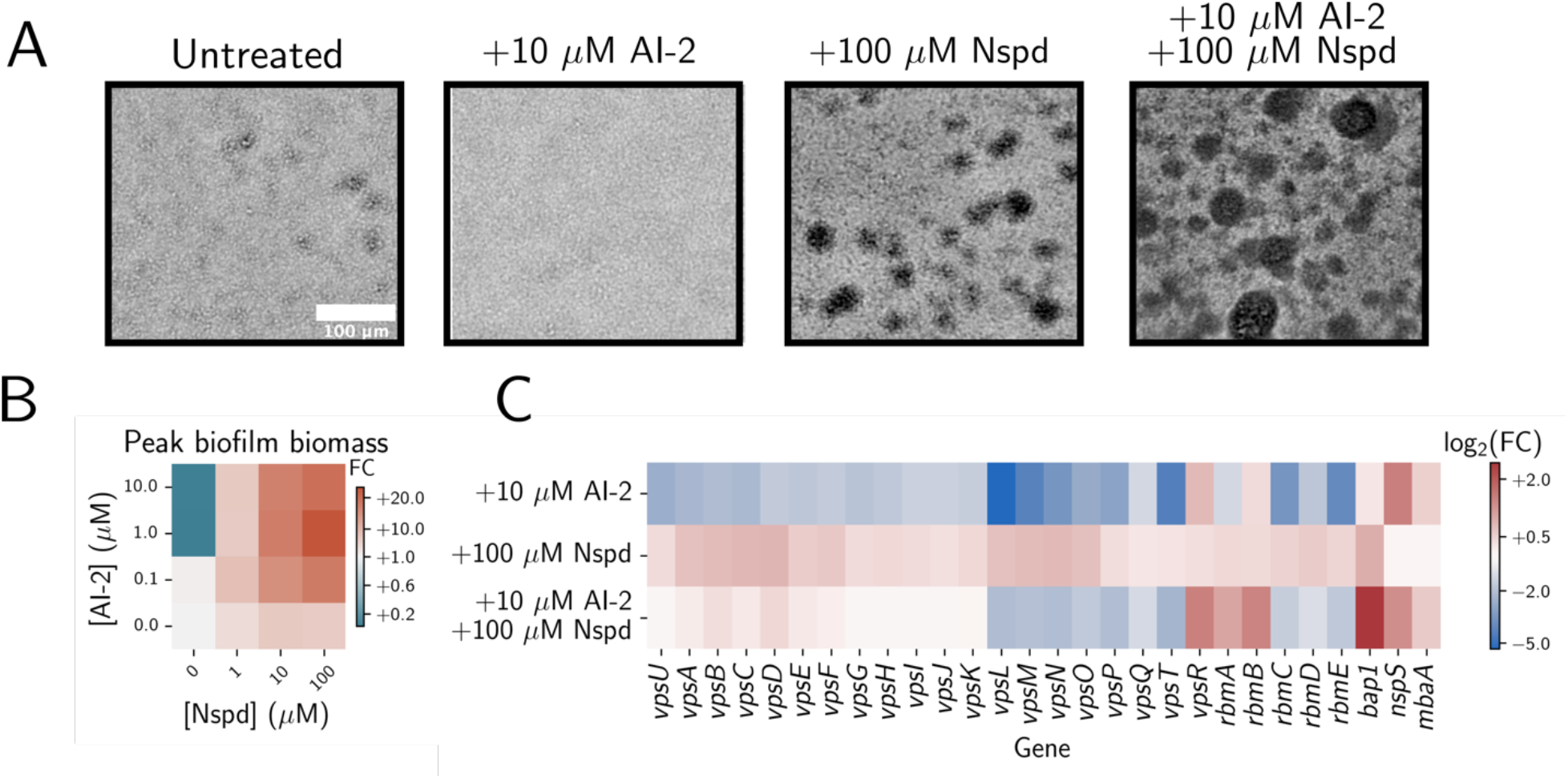
Quorum sensing promotes norspermidine-mediated biofilm formation in *V. cholerae*. (A) Representative brightfield images of biofilms produced by the *V. cholerae* AI-2-responsive strain after 14 h growth with the indicated treatments. (B) Quantitation of peak biofilm biomass for the AI-2-responsive strain grown with the indicated treatments, displayed as a heatmap. Data are normalized as fold changes relative to the untreated AI-2-responsive strain (bottom left corner). (C) Heatmap of log_2_ fold changes in biofilm gene expression in the AI-2-responsive strain grown with the indicated treatments normalized to that of the untreated strain. Samples were collected at OD_600_ = 0.1. Norspermidine, Nspd; Fold change, FC.

To define the gene expression changes underlying the quorum-sensing and norspermidine signaling synergy in *V. cholerae* biofilm formation, we conducted RNAseq in the AI-2-responsive strain under each condition shown in Fig. 2A. Treatment with AI-2 alone drove a reduction in *vps* operon, *vpsT, rbmA*, and *rbmC-E* expression, consistent with previous findings and with repression of biofilm formation (Fig. 2C) (21,24). Conversely, treatment with norspermidine caused a modest elevation in *vps* operon and *bap1* expression (Fig. 2C). Simultaneous treatment with AI-2 and norspermidine reduced *vps* operon, *vpsT*, and *rbmC-E* expression and increased *vpsR, rbmA*, and *bap1* expression. These results suggest that quorum sensing and norspermidine act synergistically to promote biofilm formation through a mechanism that decouples *vps* polysaccharide biosynthesis gene expression from expression of genes encoding the matrix proteins RbmA and Bap1.

### HapR activates *nspS-mbaA* expression at high cell density, which increases both c-di-GMP production and biofilm formation in response to norspermidine

To explore the unexpected result that quorum sensing enhances norspermidine-driven biofilm formation in *V. cholerae*, we began by measuring effects on c-di-GMP – the immediate output of the MbaA circuit in response to norspermidine. To do this, we employed a fluorescent, riboswitch-based reporter of c-di-GMP levels (25,26). Surprisingly, although provision of AI-2 alone repressed biofilm formation (Fig. 2A,B), c-di-GMP reporter output was modestly elevated in the high cell density quorum-sensing state (Fig. 3A). We considered possible roles for quorum-sensing master regulators in modulating c-di-GMP levels. It is known that the HapR high cell density master transcription factor drives c-di-GMP degradation eliminating it as a candidate (Fig. 1) (26). Thus, we suspected that the low cell density quorum-sensing master regulators – the Qrr1-4 small RNAs and/or the AphA transcription factor – could lower c-di-GMP levels at low cell density. If so, repression of the low cell density master regulators at high cell density could underpin the increase in c-di-GMP that occurs following AI-2 treatment. Indeed, AphA suppresses c-di-GMP reporter output (Fig. S2). Thus, both the low and high cell density quorum-sensing master regulators lower c-di-GMP levels, and high cell density repression of *aphA* expression explains how supplementation with AI-2 elevates c-di-GMP reporter output. Finally, consistent with our biofilm measurements, simultaneous administration of norspermidine and AI-2 drove maximal c-di-GMP reporter output (Fig. 3A).

**Fig. 3.**
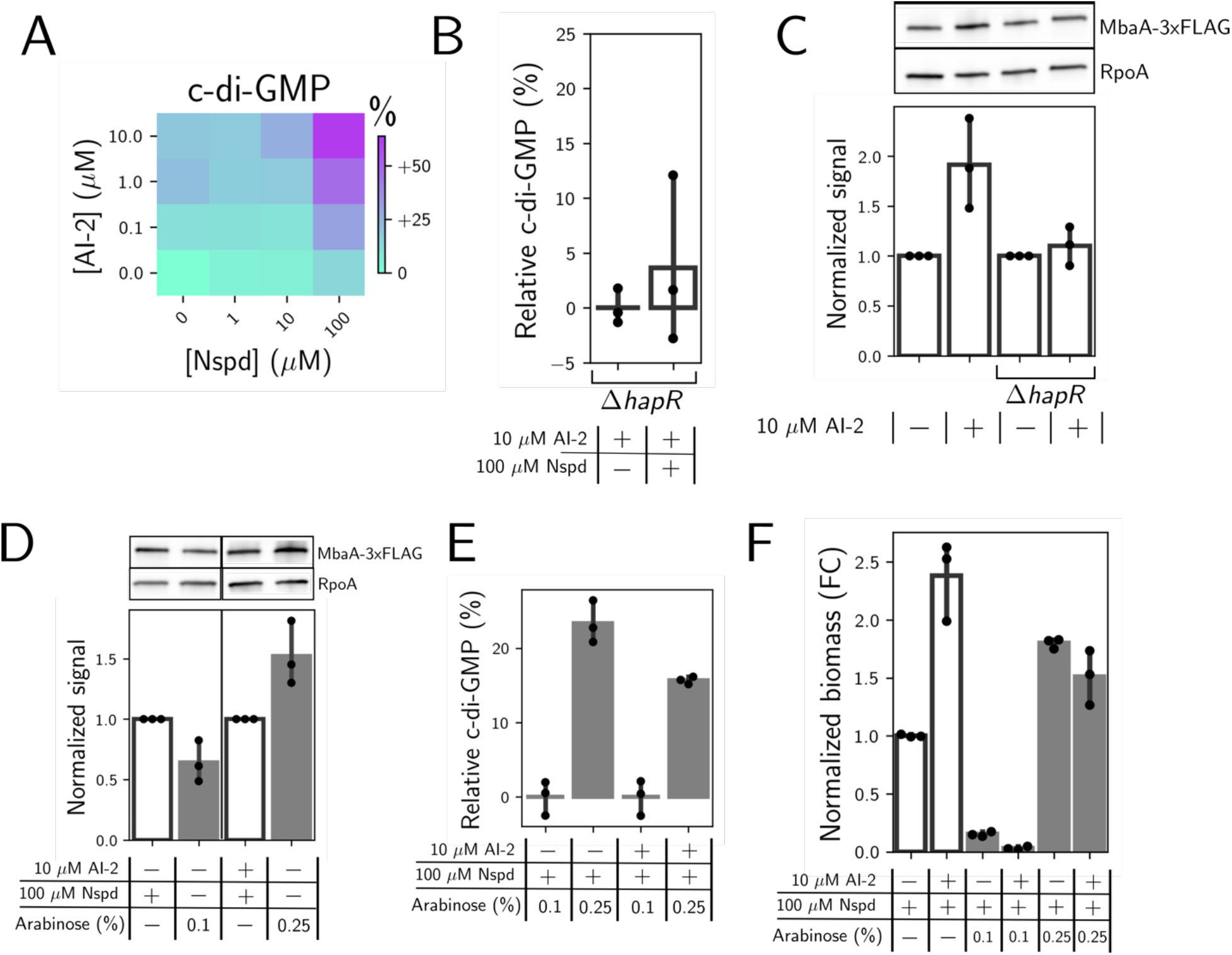
HapR-mediated activation of *nspS-mbaA* expression drives quorum-sensing and norspermidine synergy in c-di-GMP production and biofilm formation. (A) c-di-GMP reporter output in the AI-2-responsive strain following the indicated treatments, shown as a heatmap. Data are displayed as percent differences compared to the untreated strain (bottom left corner), with teal representing low and purple representing high c-di-GMP reporter output, respectively. (B) c-di-GMP reporter output in the Δ*hapR* AI-2-responsive strain following the indicated treatments. Data are normalized as percent changes relative to the Δ*hapR* AI-2-responsive strain treated with AI-2 (left bar). *N =* 3 biological replicates. (C) Top panel: western blot of MbaA-3xFLAG in the AI-2-responsive strain and the Δ*hapR* AI-2-responsive strain following the indicated treatments. Bottom panel: quantitation of MbaA-3xFLAG protein levels from the top panel. Data are normalized as fold changes relative to the AI-2 treatment in each strain and in each replicate. *N* = 3 biological replicates. (D) Top panel: western blot of MbaA-3xFLAG in the AI-2-responsive strain (1^st^ and 3^rd^ lanes) and the AI-2-responsive strain carrying *Pbad-nspS-mbaA* on the chromosome (2^nd^ and 4^th^ lanes), treated as indicated. Bottom panel: quantitation of MbaA-3xFLAG protein levels from the top panel. Data in the first and second bars are normalized to data in the first bar for each replicate. Data in the third and fourth bars are normalized to data in the third bar for each replicate. *N* = 3 biological replicates. (E) c-di-GMP reporter output in the AI-2-responsive strain carrying *Pbad-nspS-mbaA* on the chromosome. Data are normalized as percent changes relative to the mean c-di-GMP output for the 0.1% arabinose treatment for each group. *N =* 3 biological replicates. (F) Quantitation of peak biofilm biomass for the AI-2-responsive strain and the AI-2-responsive strain carrying *Pbad-nspS-mbaA* on the chromosome, treated as indicated. Images were taken at 14 h. Data are normalized as fold changes (FC) relative to the AI-2-responsive strain grown with norspermidine. *N =* 3 biological replicates. In D-F, white bars show results for the AI-2-responsive strain, and gray bars show results for the AI-2-responsive strain carrying *pbad-nspS-mbaA-3xFLAG*. Norspermidine, Nspd; Arabinose, Ara; Fold change, FC.

To explain how AI-2 supplementation could increase c-di-GMP levels when norspermidine is present, we posited that at high cell density, HapR could activate *nspS*-*mbaA* expression. Consequently, higher levels of NspS and MbaA would be produced, enabling increased synthesis of c-di-GMP and, in turn, increased biofilm formation in response to norspermidine. Data supporting this possibility are the following: First, our RNAseq results show that in the high cell density quorum-sensing regime, *nspS* and *mbaA* transcript levels are elevated (Fig. 2C). Second, a mathematical model that we previously developed to capture NspS-MbaA-mediated c-di-GMP production/degradation predicts that elevating NspS and MbaA concentrations should increase c-di-GMP in response to norspermidine (17). Third, in the *ΔhapR* AI-2-responsive strain, c-di-GMP output remained insensitive to the addition of norspermidine when AI-2 was supplied (Fig. 3B). Thus, a HapR-dependent mechanism must underlie the elevated sensitivity of the c-di-GMP reporter to norspermidine. To test our hypothesis, we measured MbaA protein levels by western blot in the AI-2-responsive strain and in the *ΔhapR* AI-2-responsive strain in the presence and absence of AI-2. We did not measure NspS, because *nspS* and *mbaA* are in an operon, and we observed that both *nspS* and *mbaA* transcript levels increased in step in the high cell density quorum-sensing signaling state (Fig. 2C) (27). Indeed, MbaA levels doubled following AI-2 supplementation, and moreover, this increase depended on HapR (Fig. 3C).

To probe whether increasing NspS-MbaA levels is sufficient to promote the observed increase in the sensitivity of c-di-GMP biosynthesis to changes in norspermidine levels, we replaced the endogenous chromosomal *nspS-mbaA* promoter with the arabinose-controlled *Pbad* promoter, and additionally, we tagged MbaA with 3xFLAG. Thus, we could synthetically modulate NspS-MbaA production by supplying arabinose, we could quantify MbaA levels by western blot, and we could track changes in c-di-GMP production. Importantly, this strategy provided the essential feature of removing quorum-sensing control of *nspS-mbaA* transcription. We identified a concentration of arabinose (0.1%) that drove MbaA production to roughly the level achieved by norspermidine treatment alone (Fig. 3D). We likewise identified a concentration of arabinose (0.25%) that produced the doubling in MbaA production that occurs following norspermidine and AI-2 co-treatment (Fig. 3D). Companion measurements of c-di-GMP reporter output showed that increasing NspS and MbaA levels drove increased c-di-GMP production (Fig. 3E) for samples grown with only norspermidine and with both norspermidine and AI-2. Consistent with this finding, increasing NspS and MbaA levels increased biofilm biomass accumulation to roughly the same extent in the presence of norspermidine alone and in the presence of both norspermidine and AI-2 (Fig. 3F). Thus, we conclude that HapR-directed activation of *nspS*-*mbaA* expression accounts for the increased sensitivity of c-di-GMP biosynthesis to norspermidine in the high cell density quorum-sensing regime. Moreover, the increased sensitivity of c-di-GMP biosynthesis to norspermidine results in elevated biofilm biomass in the high cell density and high norspermidine signaling regime.

Finally, we considered the possibility that an NspS-MbaA-independent mechanism could also contribute to the synergy between norspermidine and quorum-sensing signaling. For this analysis, we introduced the *vpvC*^W240R^ gene encoding a constitutively active diguanylate cyclase under the *Pbad* promoter onto the chromosome of the AI-2-responsive strain. This construct allowed us to ramp up intracellular c-di-GMP levels via arabinose treatment. In the high cell density quorum-sensing regime, no increase in biofilm formation occurred at any level of *vpvC*^W240R^ expression within the range tested, suggesting that quorum sensing does not generally enhance the sensitivity of biofilm formation to changes in c-di-GMP levels (Fig. S3). Rather, quorum sensing specifically enhances norspermidine-dependent biofilm formation through an NspS-MbaA-directed increase in the sensitivity of c-di-GMP biosynthesis to norspermidine.

### MbaA synthesized c-di-GMP activates VpsR

We sought to identify the downstream component responsible for transducing the AI-2-norspermidine-driven increase in c-di-GMP into the control of biofilm formation. We hypothesized that the increased c-di-GMP produced by MbaA could activate and/or increase the levels of the transcription factors VpsT and VpsR, both of which control expression of biofilm-related genes (24). Consistent with our RNAseq results, VpsT-3xFLAG and VpsR-3xFLAG levels increase following supplementation with both norspermidine and AI-2 compared to supplementation with AI-2 alone, as does the downstream matrix protein, RbmA-3xFLAG (Figs. 2C and S4). Thus, we examined the individual roles of VpsT and VpsR in controlling RbmA protein levels. Regarding VpsT: in a Δ*vpsT* AI-2-responsive strain in the high norspermidine and high quorum-sensing signaling regime, the VpsR-3xFLAG level was equivalent to that in the AI-2-responsive strain following the same treatment (Fig. S4). However, the Δ*vpsT* AI-2-responsive strain possessed lower RbmA-3xFLAG than the AI-2-responsive strain in the high norspermidine and high quorum-sensing signaling regime (Fig. S4). Regarding VpsR: In the Δ*vpsR* AI-2-responsive strain, we could not detect VpsT-3xFLAG or RbmA-3xFLAG in the high cell density and high norspermidine signaling state (Fig. S4). Together, these results suggest that VpsR regulates *vpsT* expression, but not vice versa, and both VpsR and VpsT independently regulate *rbmA* expression. Moreover, we infer that because VpsT does not regulate *vpsR* expression, activation of *vpsR* expression in the high norspermidine and high quorum-sensing signaling regime occurs through VpsR autofeedback, as shown previously (28). We conclude that in the high cell density quorum-sensing and high norspermidine signaling regime, HapR-mediated activation of *nspS*-*mbaA* increases norspermidine-driven c-di-GMP production. c-di-GMP, in turn, activates VpsR. The VpsR-c-di-GMP complex activates expression of the *vps* operons, *rbmA*, and *vpsR*. VpsR-c-di-GMP also indirectly activates these same genes via induction of *vpsT* expression and consequent VpsT-c-di-GMP-mediated transcriptional activation.

### Activation of *rbmA* expression promotes alterations in biofilm morphogenesis in the high cell density and high norspermidine signaling regime

To probe whether quorum-sensing and c-di-GMP signaling synergistically affect overall biofilm architecture, we compared the spatial characteristics of *V. cholerae* biofilms receiving no treatment, treatment with norspermidine, and treatment with both norspermidine and AI-2 using single-cell resolution microscopy. Cells in biofilms treated with both ligands resided in closer proximity to one another at the biofilm core than cells in untreated biofilms or cells in biofilms treated with norspermidine alone (Fig 4. A-C, E). These results indicate that the high norspermidine and high cell density quorum-sensing signaling state alters global biofilm architecture, leading to densification of the biofilm core.

**Fig 4.**
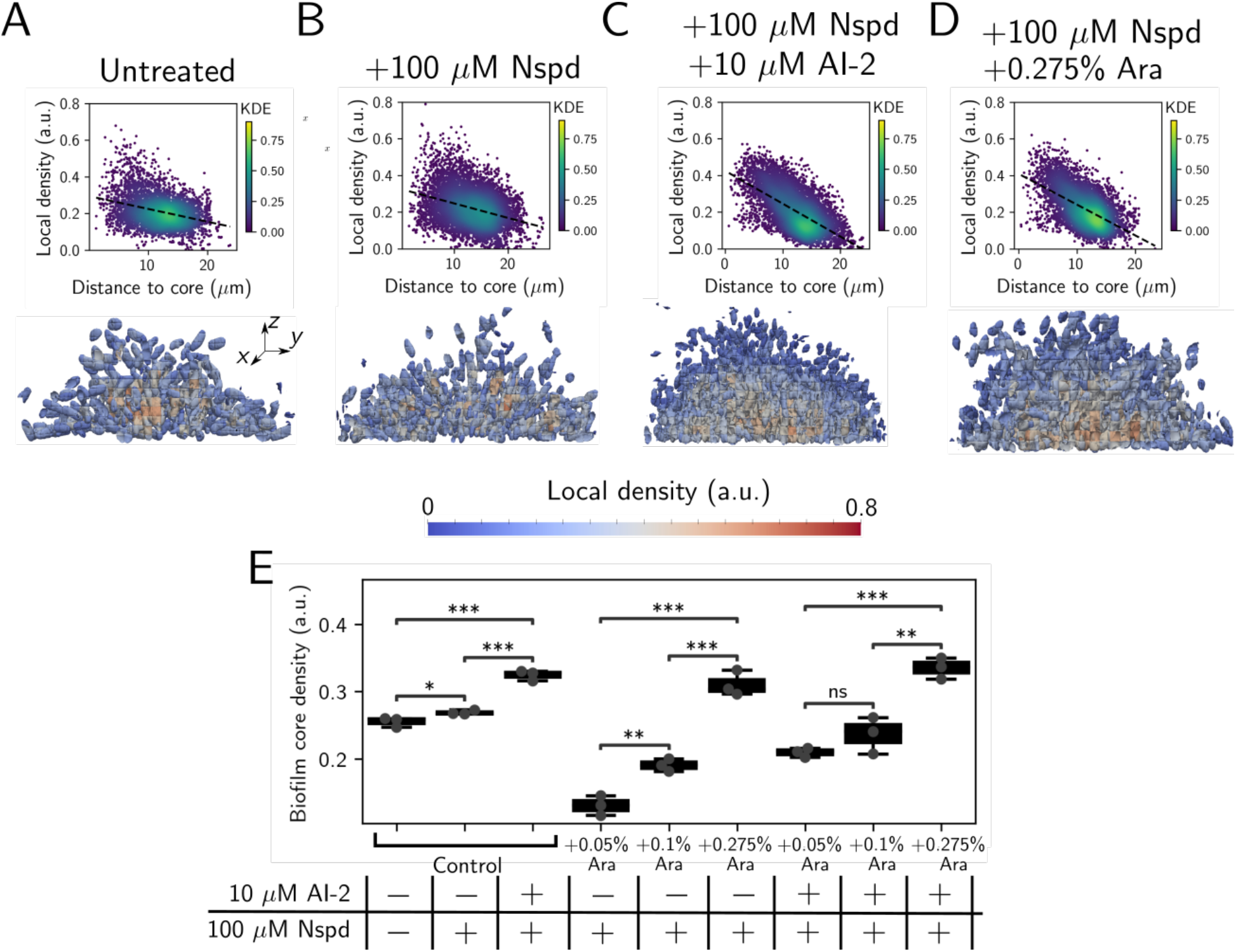
Norspermidine and quorum-sensing signaling jointly enhance biofilm formation through RbmA-mediated biofilm core densification. (A-C) (Top panels) Scatter plots showing the relationship between local biofilm cell density and distance from the biofilm core in the AI-2-responsive strain treated as indicated. (Bottom panels) Cross-sectional 3D renderings of segmented cells in biofilms ∼16 h post-inoculation, colored by local biofilm cell density and treated as in the top panels. (D) As in panels A-C for the Δ*rbmA* AI-2-responsive strain harboring chromosomal *Pbad-rbmA-3xFLAG*, treated as indicated. (E) Box plot showing the biofilm core cell density in the AI-2-responsive strain (denoted “Control”) and the Δ*rbmA Pbad-rbmA-3xFLAG* AI-2-responsive strain following the indicated treatments. Shown are the means ± standard deviations for *N =* 3 biological replicates. Unpaired t tests were performed for statistical analyses. In A-D, data points are colored by the kernel density estimate, which represents the probability density function with respect to local biofilm density and distance to the biofilm core. \*\*\*\**P* ≤ 0.0001; \*\*\**P* ≤ 0.001; \*\**P* ≤ 0.01; \**P* ≤ 0.05; ns *P* > 0.05. Norspermidine, Nspd; Arabinose, Ara; Kernel density estimate, KDE.

Obvious candidates to connect norspermidine and quorum-sensing signaling to biofilm densification are the biofilm matrix proteins Bap1 and RbmA, as expression of the genes encoding them is activated in the high cell density and high norspermidine signaling regime (Fig. 2C). Following treatment with both ligands, the Δ*bap1* strain exhibited no change in bulk biofilm biomass or biofilm core density compared to the parent AI-2-responsive strain treated with both ligands eliminating a role for Bap1 (Fig. S5A,B). By contrast, deletion of *rbmA* reduced peak biofilm biomass in the high cell density and high norspermidine signaling regime (Fig. S5A). Moreover, upon washing, biofilms formed by the Δ*rbmA* AI-2-responsive strain detached from the substrate, likely because they are fragile due to decreased cell-cell adhesion (Fig. S5C) (29,30). Synthetic induction of *rbmA* expression increased biofilm core density in a dose-dependent manner (Fig. 4E), consistent with previous results (31). Thus, ligand-driven *rbmA* upregulation is a potential mechanism that links the high norspermidine and high cell density quorum-sensing signaling regime to changes in biofilm architecture. Indeed, when we matched RbmA-3xFLAG levels in the norspermidine-treated Δ*rbmA* strain to the doubly ligand-treated parent strain using a chromosomal *Pbad-rbmA-3xFLAG* construct (using 0.275% arabinose, Figure 4D), the spatial density correlations and the biofilm core densities of the two strains became roughly equivalent (Fig. 4C-E). Thus, increased *rbmA* expression largely explains the synergistic effects of norspermidine and quorum-sensing signaling on biofilm formation.

## Discussion

In this study, we investigated the effects of simultaneously altering c-di-GMP and quorum-sensing signaling on *V. cholerae* biofilm formation. Strikingly, we found that changing c-di-GMP signaling through norspermidine supplementation had little effect on biofilm formation in the low cell density quorum-sensing signaling state but had a biofilm-promoting effect in the high cell density quorum-sensing signaling state (Fig. 2). We demonstrated that the synergy between the signaling pathways is a consequence of increased production of NspS and MbaA at high cell density. Thus, under this condition, c-di-GMP levels can increase if norspermidine is present (Fig. 3). The effect of elevated c-di-GMP levels is activation of VpsR, which we infer undergoes positive feedback and activates *rbmA* and *vps* operon gene expression both directly and indirectly via induction of *vpsT* (Figs. 2, S4). These combined changes drive the formation of larger, denser biofilms than those that form in the low cell density signaling state (Fig. 4). The major takeaway from this research is that, remarkably, quorum sensing can either promote or suppress biofilm formation, depending on the presence or absence of environmental cues that impinge on c-di-GMP signaling (Fig. 5).

**Fig. 5.**
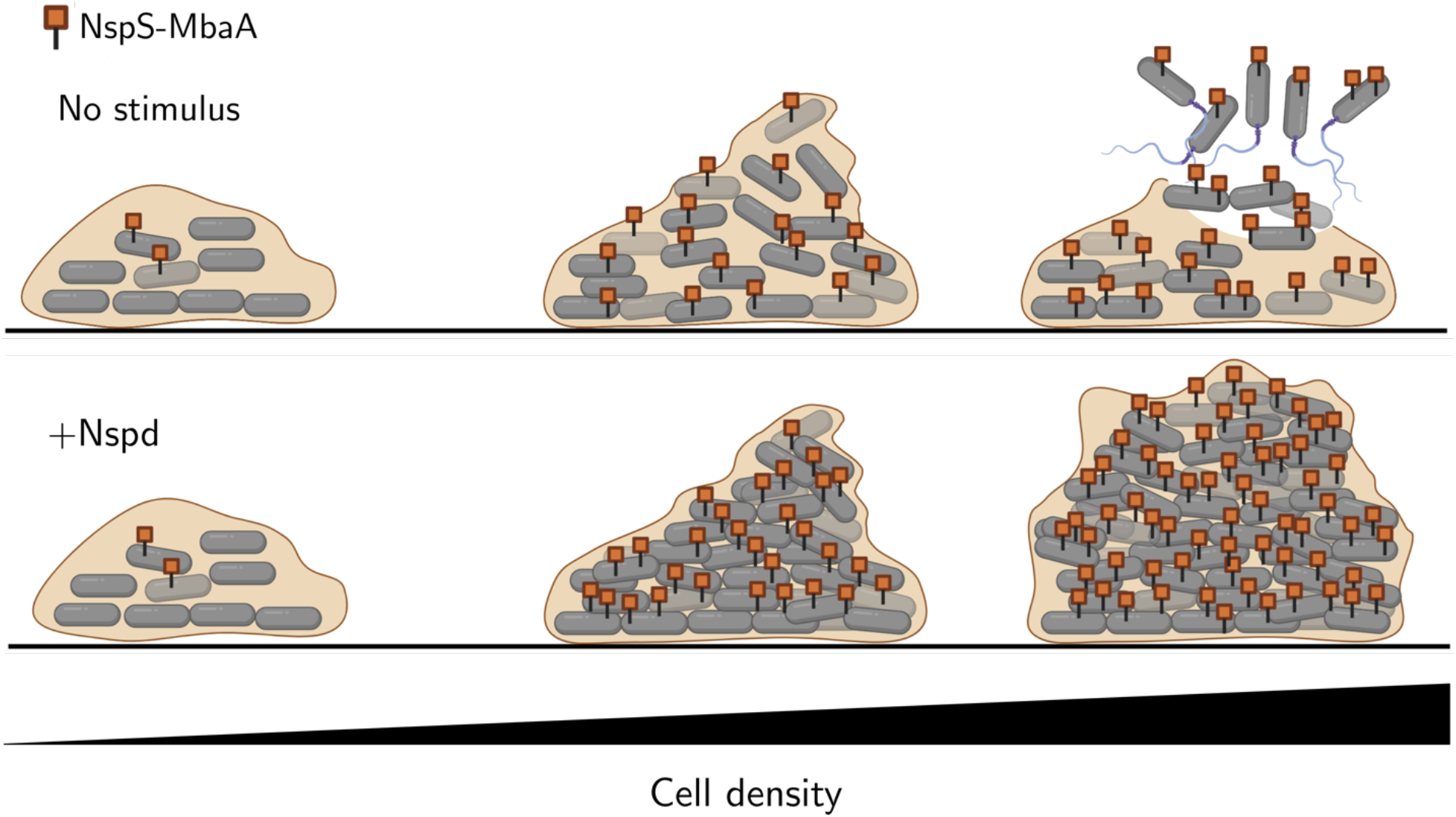
Proposed model for the integration of quorum-sensing and c-di-GMP signaling in *V. cholerae* biofilm morphogenesis. At low cell density, HapR levels are low, and consequently, NspS-MbaA levels are low, biofilm genes are expressed, and biofilms form. As the bacterial population grows and cell density increases, HapR levels rise, and HapR activates *nspS* and *mbaA* expression. (Top) At high cell density, in the absence of norspermidine, the NspS-MbaA circuit is inactive, and HapR-mediated repression of biofilm gene expression causes dispersal. (Bottom) At high cell density, in the presence of norspermidine, the NspS-MbaA pathway is activated, and high levels of the Nspd-NspS-MbaA complex produce c-di-GMP that increases biofilm gene expression, leading to biofilm expansion and densification. Norspermidine, Nspd.

Our findings imply that quorum sensing confers plasticity to the population-level decision to commit to the biofilm or the free-swimming state. In the absence of c-di-GMP-modulating signals, quorum sensing promotes the free-swimming state at high cell density, but via upregulation of c-di-GMP-metabolizing enzymes that detect environmental stimuli, quorum-sensing signaling has the potential to drive the opposite output behavior of population-level commitment to the biofilm state. It has long been known that quorum sensing controls the expression of genes encoding over a dozen c-di-GMP-metabolizing enzymes (Fig. S6) (21). However, the ramifications of this regulatory arrangement have remained mysterious prior to this work. A previously reported model for c-di-GMP and quorum-sensing integration proposed that quorum-sensing communication and detection of environmental stimuli like oxygen, polyamines, nitric oxide, etc., independently contribute to alterations in c-di-GMP levels (32). Our results show that, at least for the quorum-sensing and polyamine cues, this is not the case. Rather, the stimuli act synergistically. Testing the generality of this model remains to be performed, however, the possibility to do so is limited by the scarcity of known ligands that control diguanylate cyclase and phosphodiesterase activities.

We wonder how the results presented here might extend to other bacteria. *V. cholerae* is unusual in that individually, the high cell density quorum-sensing state and the high c-di-GMP state promote opposite biofilm phenotypes. In other bacteria, such as *Pseudomonas aeruginosa*, the high cell density quorum-sensing state and the high c-di-GMP state both independently promote biofilm formation (33,34). In *P. aeruginosa*, the prevailing model for c-di-GMP signaling and its influence on biofilm development is that c-di-GMP-metabolizing enzymes with specialized sensory functions in biofilm formation (e.g., surface sensing) are upregulated and/or activated at different points in the biofilm lifecycle, typically via two-component signal transduction pathways (34). Thus, context-dependency is a known feature of c-di-GMP signaling in *P. aeruginosa* biofilm morphogenesis, however, connections between quorum sensing and environmental stimuli that promote changes in c-di-GMP levels and biofilm formation remain uncharacterized in *P. aeruginosa*. Probing the interactions between quorum-sensing and c-di-GMP signaling in *P. aeruginosa* and other species that occupy diverse niches and that have lifestyles that differ dramatically from that of *V. cholerae* could deliver a unified picture of how the coordination of sensory signaling systems is linked to the ecological and evolutionary roles that biofilms play across the bacterial domain.

## Methods

### Bacterial strains, reagents, reporters, and western blotting procedures

The *V. cholerae* strain used in this study was O1 El Tor biotype C6706str2. Antibiotics were used at the following concentrations: polymyxin B, 50 µg/mL; kanamycin, 50 µg/mL; spectinomycin, 200 µg/mL; chloramphenicol, 1 µg/mL; and gentamicin, 5 µg/mL. Strains were propagated at 30° C in liquid lysogeny broth (LB) with shaking or LB containing 1.5% agar for plates. Strains used for reporter assays, imaging assays, and RNA isolation were grown in M9 minimal medium supplemented with 0.5% dextrose, 0.5% casamino acids, and 0.1 mM boric acid. AI-2 (*S*-2-methyl-2,3,3,4-tetrahydroxytetrahydrofuran-borate) and the CqsS agonist 1-ethyl-*N*-{[4-(propan-2-yl)phenyl]methyl}-1*H*-tetrazol-5-amine were synthesized as described previously (35–38). Norspermidine (Millipore Sigma, I1006-100G-A), arabinose (Millipore Sigma, W325501), AI-2, and the CqsS agonist were added at the concentrations designated in the figures or figure legends at the initiation of the assay. c-di-GMP was measured as described previously (17,26). Western blots for MbaA-3xFLAG, VpsT-3xFLAG, VpsR-3xFLAG, and RbmA-3xFLAG were performed as described previously, using a monoclonal anti-FLAG-peroxidase antibody (Millipore Sigma, #A8592; Danvers, MA, USA). RpoA served as the loading control and it was detected using an anti-*Escherichia coli* RNA polymerase α primary antibody (Biolegend, #663104) and an anti-mouse IgG HRP conjugate secondary antibody (Promega, #W4021) (13). For strains carrying VpsT-3xFLAG, RbmA-3xFLAG, or VpsR-3xFLAG, prior to application of the anti-RpoA antibody, the anti-FLAG-peroxidase antibody was stripped from the membranes by incubation at 25° C in stripping buffer (15 g/L glycine, 1 g/L SDS, 10 mL/L Tween-20, diluted in water, buffered to pH = 2.2) for 15 min, followed by a second incubation with stripping buffer for 10 min, followed by two 10 min incubations in PBS, and finally two 5 min incubations in PBST.

### DNA manipulation and strain construction

Modifications to the *V. cholerae* genome were generated by replacing genomic DNA with linear DNA introduced by natural transformation as described previously (13,39,40). PCR and Sanger sequencing (Genewiz) were used to verify genetic alterations. See S1 Table for primers and g-blocks (IDT) and S2 Table for a list of strains used in this study. Constructs driven by the *Pbad* promoter were introduced at the neutral locus *vc1807*. The *Pbad-nspS-mbaA* construct was produced by replacing the native *nspS* promoter with *Pbad*.

### Microscopy Analyses

Measurements of biofilm biomass were made as described previously (13) using bright field microscopy with minimal modifications. In brief, single-plane images were acquired at 30 min intervals on a Biotek Cytation 7 multimodal plate reader using an air immersion 20x objective lens (Olympus, PL FL; NA: 0.45) with static incubation at 30° C. Analyses were performed using FIJI software (Version 1.53c). Images in the time-series were smoothed using a Gaussian filter (σ = 10), followed by segmentation using an intensity threshold. The total amount of light attenuated in each image after segmentation was summed to yield the biofilm biomass for the corresponding time point.

For high resolution images of cells in biofilms (Figs. 4 and S5), samples were fixed by treatment with 3.7% formaldehyde (Avantor, MFCD00003274) in PBS for 10 min. To terminate fixation, samples were washed five times with PBS. Cells were subsequently stained with 1 µg/mL 4′,6-diamidino-2-phenylindole (DAPI) in PBS for 30 min at 25° C. Single-cell resolution images of fixed samples were acquired using a DMI8 Leica SP-8 point scanning confocal microscope (Leica, Wetzlar, Germany) equipped with a 63x water immersion objective (Leica, HC PL APO CS2; NA: 1.20). The excitation light source was a 405 nm diode laser and emitted light was detected by a GaAsP spectral detector (Leica, HyD SP). Cell segmentation and biofilm parameter calculations were performed using BiofilmQ (parameters Architecture_LocalDensity and Distance_ToBiofilm CenterAtSubstrate) (31). All plots were generated using Python 3. Figures were assembled in Inkscape (41,42).

### RNA isolation and sequencing

Overnight cultures of the *V. cholerae* AI-2-responsive strain, grown in biological triplicate, were diluted to OD_600_ ∼ 0.001 in 5 mL of M9 medium. The subcultured cells were grown at 30° C with shaking in the presence of the designated polyamine and/or AI-2 treatment to OD_600_ = 0.1. Cells were harvested by centrifugation for 10 min at 4,000 RPM and resuspended in RNAprotect (Qiagen). RNA was isolated using the RNeasy mini kit (Qiagen), remaining DNA was digested using the TURBO DNA-free kit (Invitrogen), and the concentration and purity of RNA were measured using a NanoDrop instrument (Thermo). Samples were flash frozen in liquid nitrogen and stored at −80° C until they were shipped on dry ice to SeqCenter (https://www.seqcenter.com/rna-sequencing/). The 12 million paired-end reads option and the intermediate analysis package were selected for each sample. Quality control and adapter trimming were performed with bcl2fastq (Illumina), while read mapping was performed with HISAT2 (43). Read quantitation was performed using the Subread’s featureCounts (44) functionality, and subsequently, counts were loaded into R (R Core Team) and normalized using the edgeR (45) Trimmed Mean of M values (TMM) algorithm. Values were converted to counts per million (cpm), and differential expression analyses were performed using the edgeR Quasi-Linear F-Test (qlfTest) functionality against treatment groups, as indicated. The results, presented in Fig. 2C, were plotted using Python 3 (41).

## Acknowledgements and Funding

The authors thank members of the Bassler group for insightful discussions. This work was supported by the Howard Hughes Medical Institute, NSF grant MCB-2043238, NIH grant 2R37GM065859 (B.L.B.), and NIH grant 1K99AI158939 (A.A.B). The funders had no role in study design, data collection and analysis, decision to publish, or preparation of the manuscript.

---

**Fig. S1.**
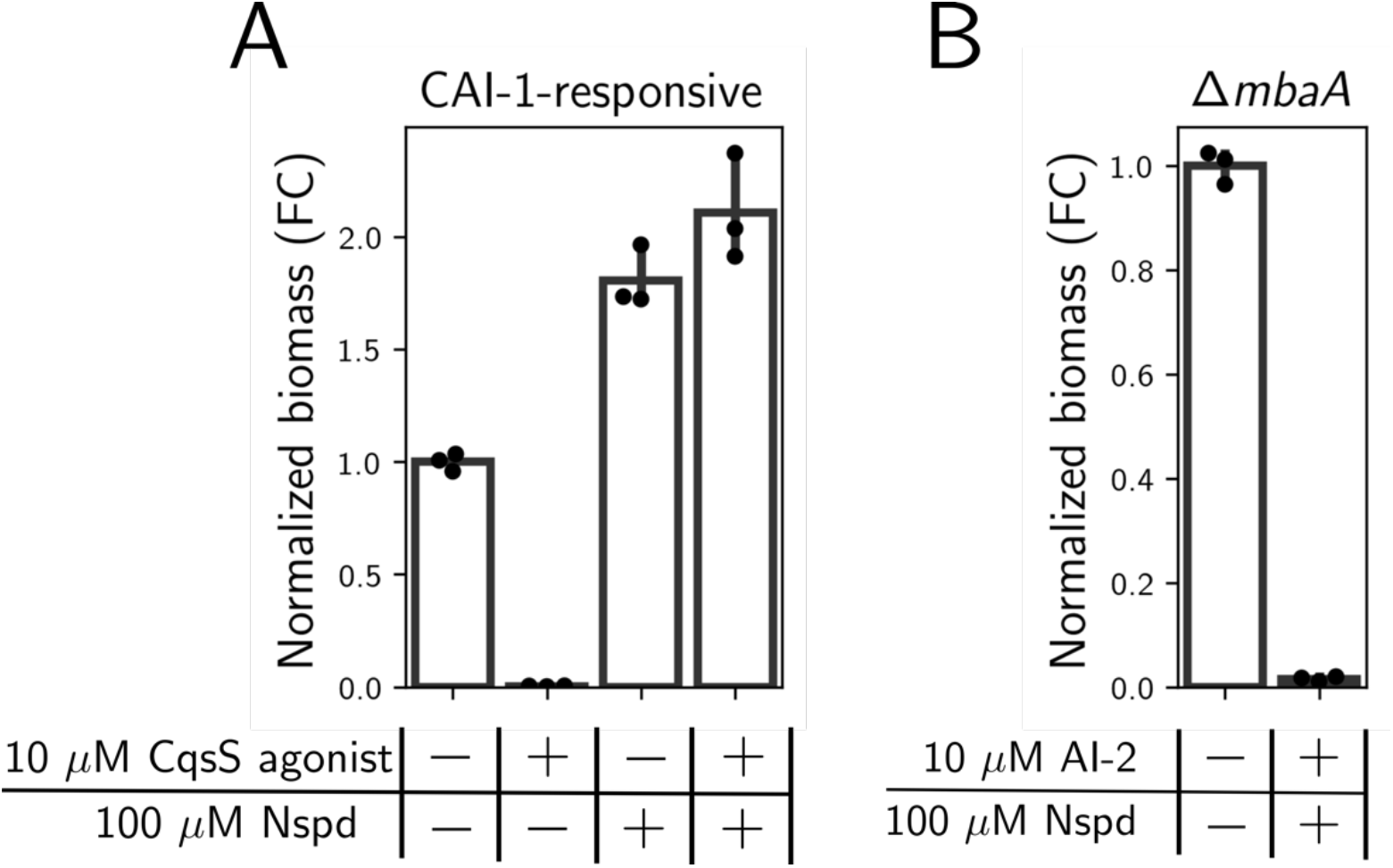
Synergy between norspermidine and AI-2 treatments is mediated by the NspS-MbaA and quorum-sensing pathways. (A) Peak biofilm biomass, measured by quantitative brightfield imaging of biofilms produced by the CAI-1-responsive strain grown with the indicated treatments. (B) Peak biofilm biomass produced by the Δ*mbaA* AI-2-responsive strain, grown with the indicated treatments. Data are normalized as fold changes relative to the untreated strains. *N =* 3 biological replicates. Norspermidine, Nspd; Fold change, FC.

**Fig. S2.**
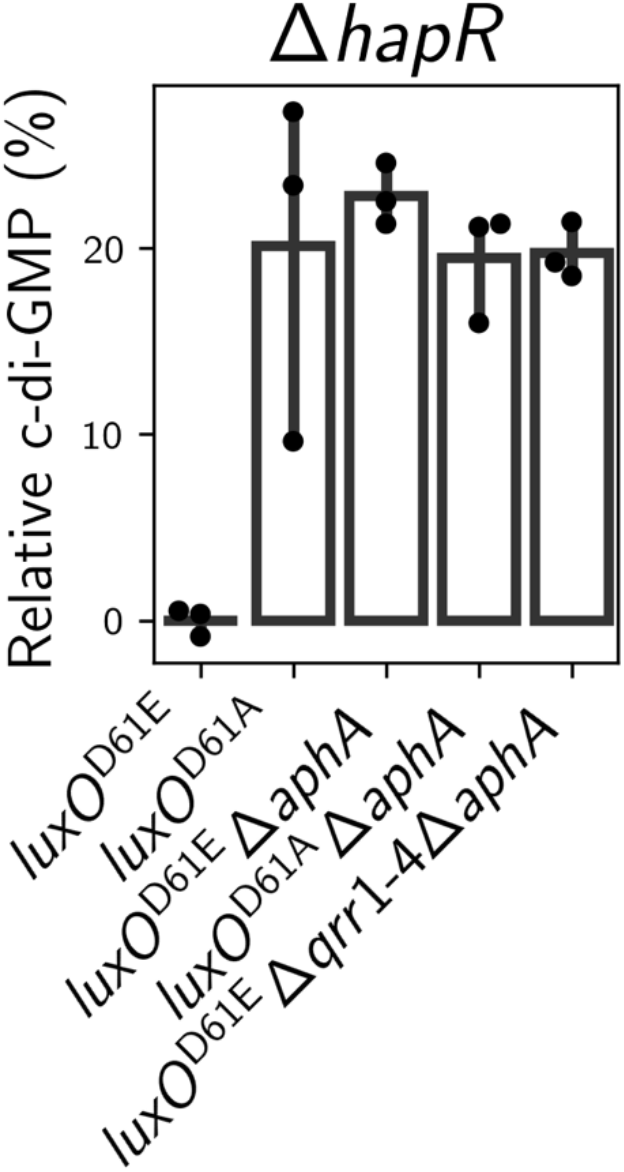
AphA activity lowers cytoplasmic c-di-GMP levels. c-di-GMP reporter output in the indicated *V. cholerae* strains. Data are normalized as percent changes relative to the c-di-GMP produced by the *luxO*^D61E^ Δ*hapR* strain. The LuxO^D61E^ mutant protein mimics LuxO∼P, thus locking *V. cholerae* in the low cell density quorum-sensing state. The LuxO^D61A^ mutant protein is incapable of phosphorylation, thus locking *V. cholerae* in the high cell density quorum-sensing state.

**Fig. S3.**
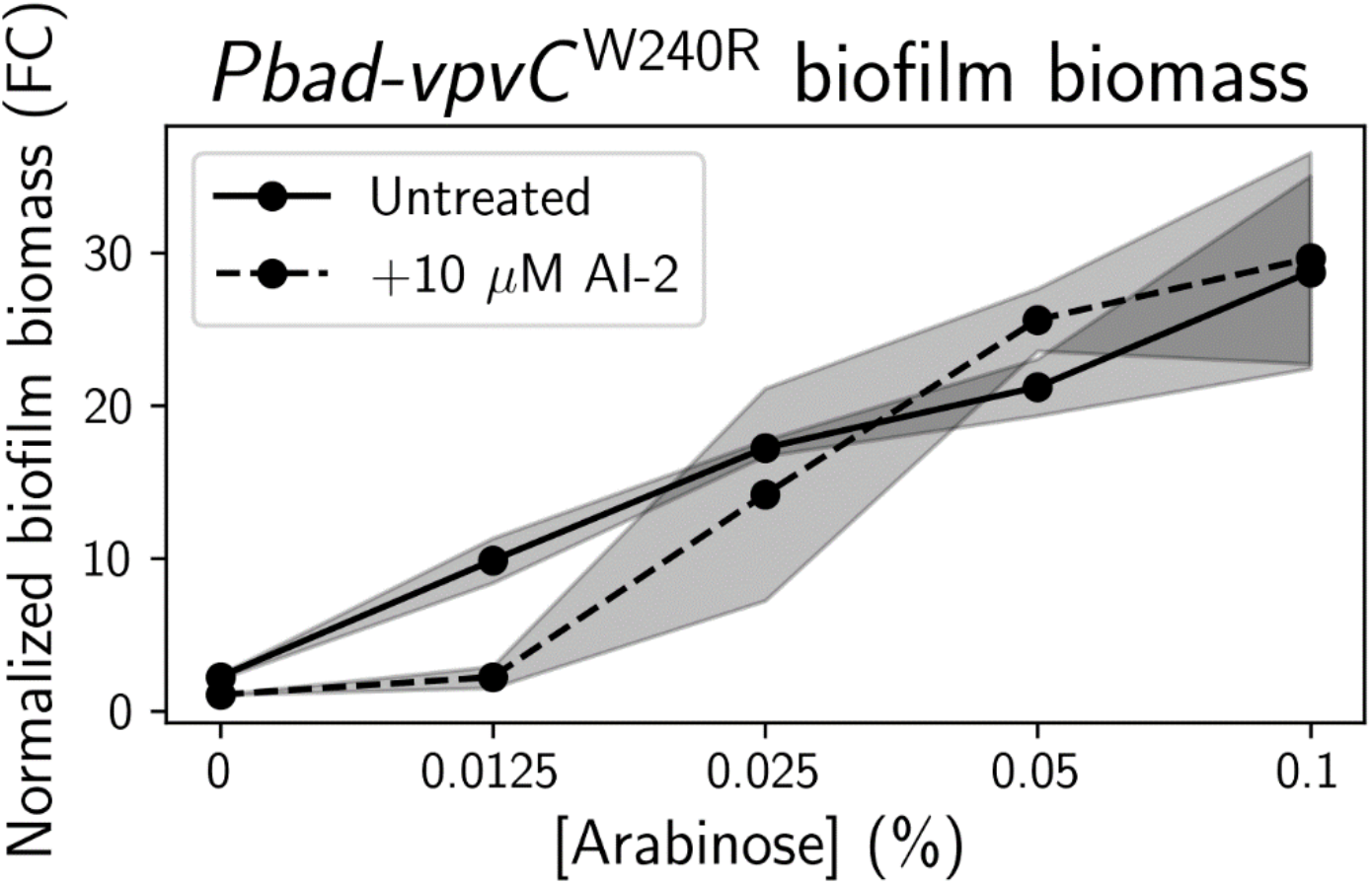
Quorum sensing does not generally alter the sensitivity of biofilms to changes in cytoplasmic c-di-GMP levels. Peak biofilm biomass in the AI-2-responsive strain carrying chromosomal *Pbad*-*vpvC*^W240R^, grown with the indicated treatments. Data are normalized as fold changes relative to the untreated strain at 0% arabinose.

**Fig. S4.**
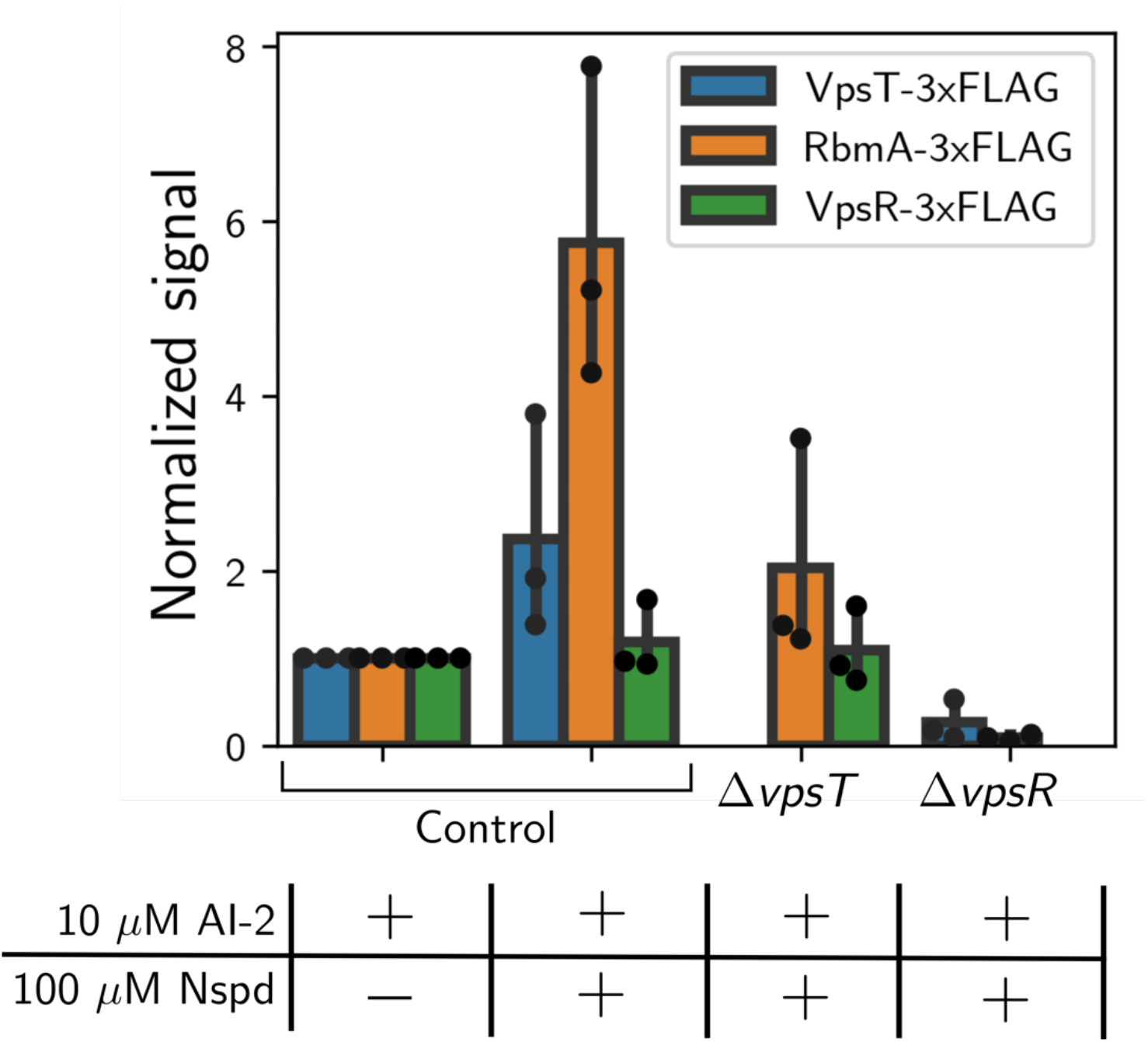
VpsR activates *vpsT* and *rbmA* expression in the high cell density and high norspermidine signaling regime. Quantitation of VpsT-3xFLAG, RbmA-3xFLAG, and VpsR-3xFLAG protein levels from western blots performed on the indicated strains with the indicated treatments. The AI-2-responsive strain is designated Control. Data are normalized to protein levels in the AI-2 treatment condition (left set of bars). Norspermidine, Nspd.

**Fig. S5.**
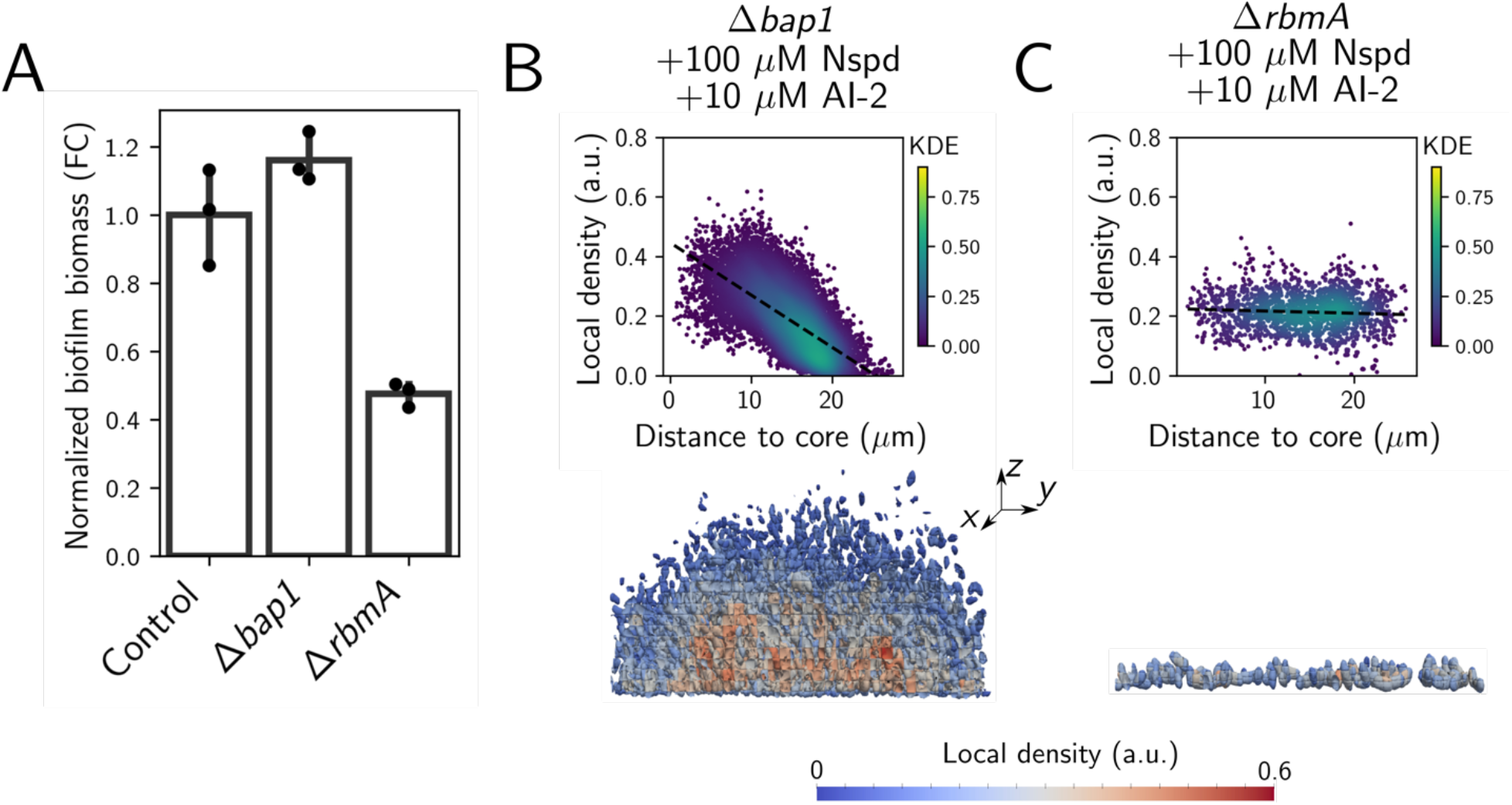
Activation of *rbmA* expression but not *bap1* expression drives changes in biofilm formation in the high cell density and high norspermidine signaling regime. (A) Peak biofilm biomass, measured by quantitative brightfield imaging of biofilms produced by the AI-2-responsive strain (designated Control), the Δ*bap1* AI-2-responsive strain, and the Δ*rbmA* AI-2-responsive strain, each treated with both norspermidine and AI-2. (B-C) (Top panels) Scatter plots showing the relationship between local biofilm cell density and distance from the biofilm core. (Bottom panels) Cross-sectional 3D renderings of segmented cells in biofilms ∼16 h post-inoculation, colored by local biofilm cell density. (B) In the Δ*bap1* AI-2-responsive strain treated with norspermidine and AI-2. (C) In the Δ*rbmA* AI-2-responsive strain treated with norspermidine and AI-2. Norspermidine, Nspd; Kernel density estimate, KDE.

**Fig. S6.**
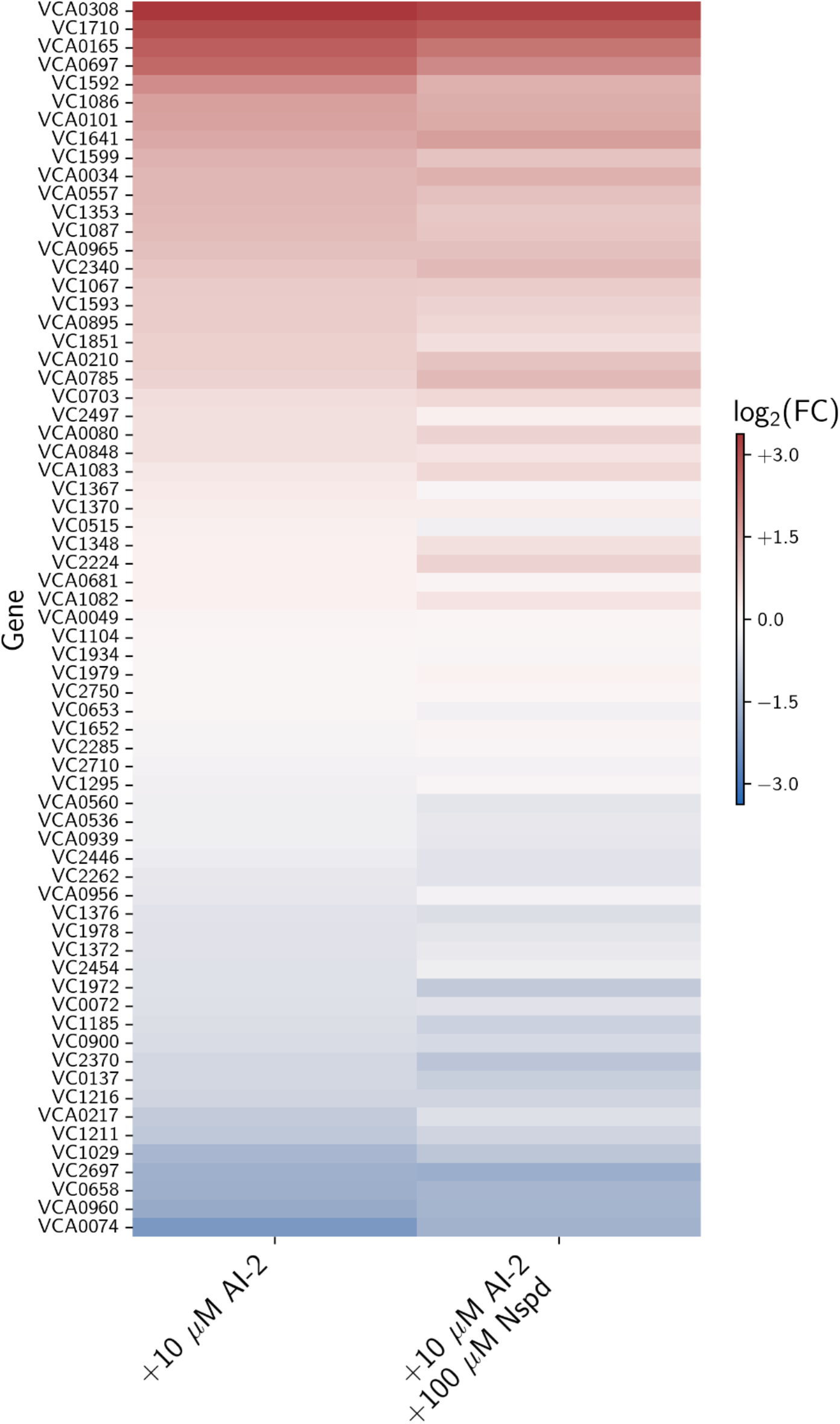
Transcriptional regulation of genes encoding c-di-GMP metabolizing enzymes by the high cell density quorum-sensing state. Heatmap of log_2_ fold changes in the expression of genes encoding diguanylate cyclases and phosphodiesterases in the AI-2-responsive strain grown with the indicated treatments normalized to transcript levels in the untreated strain. Samples were collected at OD_600_ = 0.1. Norspermidine, Nspd; Fold change, FC.

